# BicD and MAP7 collaborate to activate homodimeric *Drosophila* kinesin-1 by complementary mechanisms

**DOI:** 10.1101/2025.01.11.632512

**Authors:** M Yusuf Ali, Hailong Lu, Patricia M. Fagnant, Jill E. Macfarlane, Kathleen M. Trybus

## Abstract

The folded auto-inhibited state of kinesin-1 is stabilized by multiple weak interactions and binds weakly to microtubules. Here we investigate the extent to which homodimeric *Drosophila* kinesin-1 lacking light chains is activated by the dynein activating adaptor *Drosophila* BicD. We show that one or two kinesins can bind to the central region of BicD (CC2), a region distinct from that which binds dynein-dynactin (CC1) and cargo-adaptor proteins (CC3). Kinesin light chain significantly reduces the amount of kinesin bound to BicD and thus regulates this interaction. Binding of kinesin to BicD increases the number of motors bound to the microtubule, the fraction moving processively and the run length, suggesting that BicD relieves kinesin auto-inhibition. In contrast, microtubule-associated protein 7 (MAP7) has minimal impact on the percentage of motors moving processively but enhances both kinesin-1 recruitment to microtubules and run length. BicD relieves auto-inhibition of kinesin, while MAP7 enables activated motors to engage productively with microtubules. When BicD and MAP7 are combined, the most robust activation of kinesin-1 occurs, highlighting the crosstalk between adaptors and microtubule associated proteins in regulating transport. These observations imply that when both dynein and kinesin-1 are simultaneously bound to BicD, the direction the complex moves on MTs will be influenced by MAP7 and the number of bound kinesins.

## Introduction

The BICD family of proteins is a well-studied class of adaptors that activates the dynein-dynactin complex. BicD was first identified in *Drosophila*, where mutations resulted in an abnormal two-tailed or “bicaudal” phenotype in early fly embryos due to mis-localization of anterior and posterior mRNA polarity determinants in the oocyte (1). Mammals have two orthologs called BicD1 and BicD2, as well as two shorter related proteins called BicDR1 and BicDR2 (reviewed in (2)). BicD is a coiled-coil homodimer with three regions that have been traditionally referred to as coiled-coil 1, 2 and 3 (CC1, CC2 and CC3). The N-terminal CC1 is sufficient to recruit and activate dynein-dynactin for processive motion on microtubules (MTs) (3, 4). Like many dynein activating adaptors, BicD contains two conserved motifs: the CC1 box that interacts with the dynein light-intermediate chain and a Spindly motif that binds the pointed end of dynactin (5, 6). The C-terminal portion, CC3, interacts with adaptor proteins that specify the attached cargo. In *Drosophila*, GTP-bound Rab6 on Golgi vesicles and mRNA bound to the mRNA binding-protein Egalitarian are the best-known cargo adaptor/cargo pairs (7–9). The central region of mammalian BicD2 was shown to bind kinesin-1 (10), and kinesin was also implicated in BicD mediated motion of lipid droplets during *Drosophila* embryogenesis (11). Thus BicD has the ability to recruit both minus- and plus-end directed motors to transport cargo bidirectionally, but the effect of BicD on kinesin activation has not been well-studied.

A key feature of BicD is its ability to form an auto-inhibited conformation in which CC3 interacts with CC1. Binding of cargo to BicD drives dynein recruitment in *Drosophila* (9). Subsequent *in vitro* reconstitution experiments using purified proteins showed that binding of Egalitarian and mRNA to BicD allows recruitment and activation of dynein-dynactin by relieving the auto-inhibited state of BicD (12, 13). Electron microscopy of *Drosophila* BicD showed a looped structure formed by CC3 bending back and binding to CC1 that was disrupted only by the addition of both Egl and mRNA (13). Images reminiscent of this looped structure were also recently observed in mammalian BicD2 (14, 15). Whether the auto-inhibited state of BicD also gates access to kinesin is not known.

Both dynein and kinesin exist in auto-inhibited and activated states. The auto-inhibited state of dynein forms a structure known as the phi particle. Inhibition of microtubule (MT) binding occurs because the coiled-coil stalk regions are crossed and trapped in a registry with low affinity for the MT (16). The activation mechanism by various adaptors has been studied structurally (17, 18) and involves changes to the dynein tail that force the motor domains to adopt a parallel conformation favorable for processive motion (17, 19). Another level of regulation involves Lis1, a co-factor that binds to the dynein motor domain and promotes a conformation that is favorable for assembling an active tripartite complex with dynactin and an activating adaptor (reviewed in (20)).

The mechanism of auto-inhibition of kinesin-1 has been studied for far longer than that of dynein. Interactions between the motor domain and the globular C-terminal tail that would restrict motor domain motion was a key feature of the kinesin inhibited state, made possible via a flexible hinge region in the stalk (21–26). Remarkably, recent studies using computational modeling, crosslinking and electron microscopy showed that the hinge originally thought to allow a head-tail interaction (called “hinge-2”) is not the region where the kinesin molecule folds in half (27, 28). A newly defined “elbow” located C-terminal to hinge-2 allows formation of the inhibited molecule, which is stabilized by multiple weak interactions along the stalk, and by kinesin light chain (KLC) interactions with the heavy chain (KHC).

A popular hypothesis is that kinesin auto-inhibition is relieved by binding to cargo adaptors, a mechanism that would preclude wasting ATP hydrolysis unless cargo is coupled for transport. Recent single-molecule studies were used to investigate how the auto-inhibited mammalian heterotetrameric kinesin-1-light chain complex is functionally activated by nesprin-4, an adaptor that binds to the KLC (29). Even low concentrations of nesprin-4 increased the number of motors moving processively, but much higher nesprin-4 concentrations were needed to increase recruitment of motors to the MT. When both nesprin-4 and MAP7 were combined with heterotetrameric kinesin-1, the most robust processive motion of kinesin was observed, leading the authors to suggest that cargo adaptors and MAP7 act synergistically to activate robust mobility of kinesin (29). Previous cellular studies have implicated the microtubule-associated-protein-7 (MAP7) as a key co-factor for optimal kinesin function (30–33). *In vitro* studies showed that MAP7 interacts with both MTs and kinesin, recruits kinesin to MTs, and enhances run length and binding frequency on MTs (34, 35).

These studies raise the question of how full activation of kinesin-1 is achieved. Does the level of activation vary depending on whether kinesin is linked to its cargo via its heavy chain or light chain? Do different adaptors activate kinesin to different degrees? Is MAP7 required in all cases? Here we contribute to our understanding of kinesin-1 activation by investigating the effect of kinesin heavy chain binding to the activating adaptor BicD, which also recruits dynein-dynactin. Our results suggest that the combination of BicD and MAP7 produces the most active kinesin, consistent with the synergistic model favored by McKenney and colleagues (29). Binding of kinesin to BicD mainly relieves auto-inhibition and thus leads to more recruitment to the MT, while MAP7 predominantly enhances recruitment to the MT of already active molecules, and does not contribute significantly to relieving auto-inhibition per se. While relief of auto-inhibition and recruitment to the MT can both be considered “activation”, the two mechanisms fundamentally differ, allowing an interplay between the two mechanisms to allow fine-tuning of kinesin activity.

## Results

### Kinesin-1 heavy chain binds to the central portion of BicD

We used single-molecule techniques to identify where homodimeric *Drosophila* kinesin-1 (no bound light chains; hereafter kinesin) binds to the dynein activating-adaptor *Drosophila* BicD (hereafter BicD). A series of truncated BicD constructs were expressed and purified to identify the kinesin binding site (Fig. 1A). Kinesin lacking the hinge 2 region (kinesin^ΔH2^), which diminishes formation of the auto-inhibited state (24), was used to quantify kinesin-BicD complex formation. The general strategy was to label the purified BicD construct with a red streptavidin Qdot (emitting at 655 nm) at its N-terminus, with kinesin^ΔH2^ remaining unlabeled. Kinesin-BicD complexes moving on microtubules (MTs) were observed using total internal reflection fluorescence (TIRF) microscopy and quantified (Fig. 1B). Kinesin^ΔH2^ binds weakly to full-length BicD, which exists in an auto-inhibited looped conformation that we previously showed prevented dynein-dynactin binding (13). Neither the BicD^CC1^ construct that binds dynein-dynactin (3, 4), nor the BicD^CC3^ construct (amino acids 536-782) that binds to cargo-adaptor proteins (7–9), interacted significantly with kinesin^ΔH2^. The remaining four BicD truncation constructs (BicD^CC2^, BicD^318-658^, BicD^CC2-CC3^, and BicD^437-782^) exhibited higher numbers of moving complexes, defining a binding region for kinesin in the CC2 central portion of BicD (amino acids 437-535)(dashed red rectangle, Fig. 1A). The BicD^CC2-CC3^ construct (aa 318-782) was selected for further experiments because it showed the highest average number of moving complexes when complexed with kinesin^ΔH2^.

**Figure 1.**
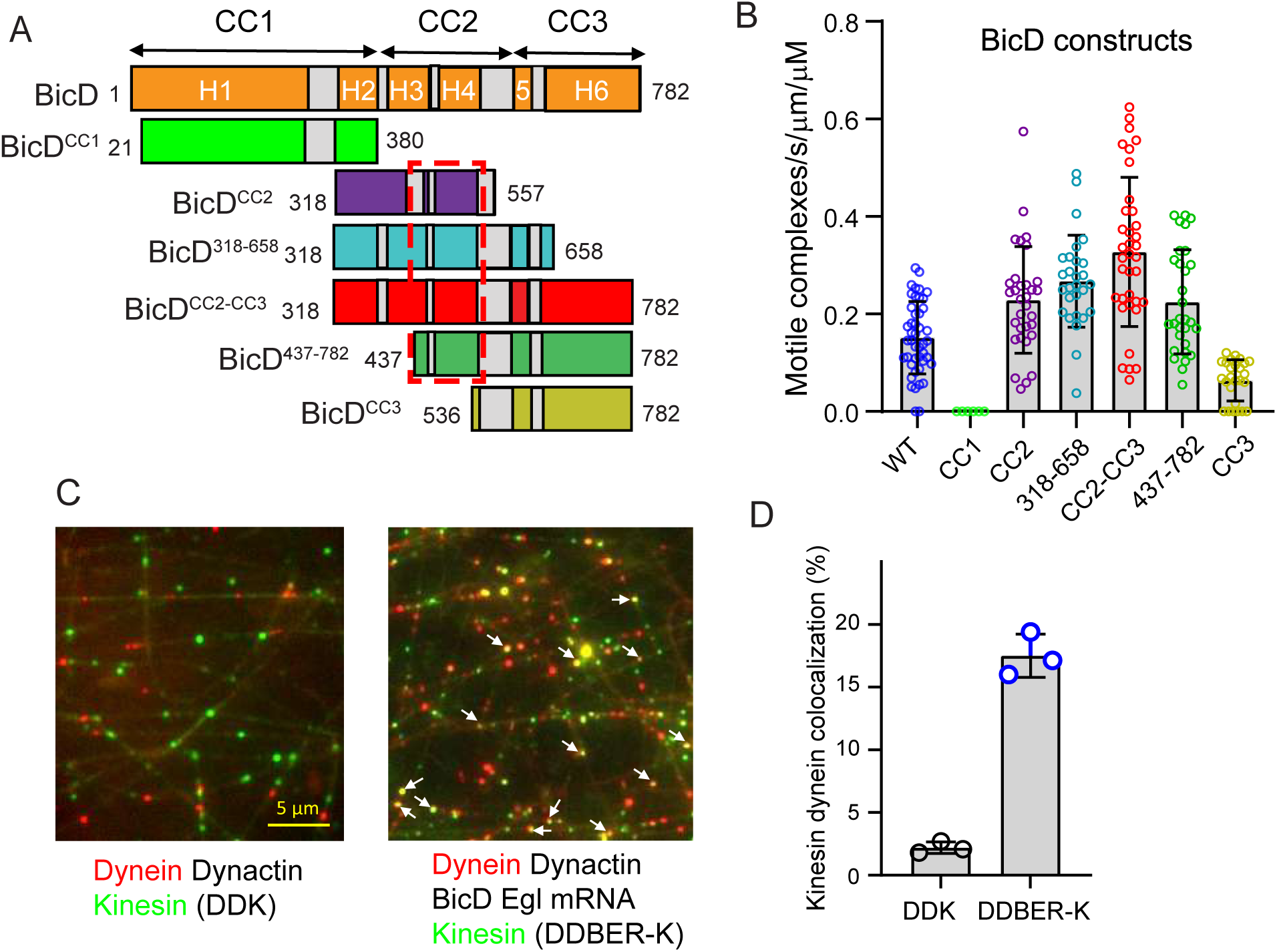
Kinesin-1 heavy chain binds to the central region of BicD. ***A***, Schematic of the series of BicD constructs used to define the kinesin binding site. Regions predicted by AlphaFold to be helical are labeled H1-H6, and the three coiled-coil domains of BicD indicated (CC1, CC2 and CC3). The red dashed boxed (amino acids 437-535) delineates the region common to truncated constructs that bind kinesin^ΔH2^ and move on microtubules (MTs) with an average frequency exceeding 0.06/sec/µm microtubule length/µM kinesin concentration. ***B***, Binding frequency of moving kinesin^ΔH2^-BicD complexes on MTs for various BicD constructs. Moving complexes were counted from n=26-45 MTs per construct. N=2 experiments. ***C***, (*right panel*) Dynein (labeled with a red Qdot) and kinesin^ΔH2^ (labeled with a green Qdot) can simultaneously bind to BicD (yellow spots, arrows) in the presence of BicD, Egalitarian (Egl) and mRNA. (*left panel*) Colocalization of dynein and kinesin was infrequently observed in the absence of BicD, Egl and mRNA. ***D***, Quantification of kinesin and dynein co-localization with dynein-dynactin and kinesin (DDK) versus dynein-dynactin, BicD, Egl, mRNA and kinesin (DDBER-K). Mean ± SD. N=2 experiments, n= 29 MTs. Unpaired t test with Welch’s correction, p<0.0001.

The above results imply that dynein, kinesin, and an adaptor protein can simultaneously bind to BicD. To directly confirm simultaneous motor binding, we added kinesin^ΔH2^ to a previously characterized reconstituted complex composed of dynein-dynactin, BicD, Egalitarian and K10 mRNA (13). Yellow complexes were observed bound to MTs when dynein was labeled with a red Qdot and kinesin with a green Qdot, indicating that both motors can simultaneously bind to full-length BicD that has been released from its auto-inhibited state by Egalitarian and K10 mRNA (Fig. 1*C*, *right panel* and 1*D*). Control experiments lacking BicD, Egalitarian and mRNA showed significantly fewer yellow complexes (Fig. 1*C*, *left panel and* 1*D*). These experiments show that dynein and kinesin can bind simultaneously to BicD, supporting the idea that these proteins function together in intracellular transport processes.

### The kinesin light chain decreases interaction between kinesin and BicD

To investigate how the kinesin light chain (KLC) impacts the interaction between kinesin and BicD, we performed experiments using kinesin constructs with or without KLC (kinesin^KLC^ or kinesin, respectively). Biotin tags at the C-terminus of the kinesin heavy chain and at the N-terminus of BicD^CC2-CC3^ were used for labeling with either red or green streptavidin-Qdots. With homodimeric kinesin heavy chain, 17% of the visualized Qdots were yellow indicating complex formation with BicD. In contrast, heterotetrameric kinesin^KLC^ only exhibited 3.7% yellow complexes (Fig. 2A). The most straightforward interpretation of these findings is that BicD and KLC compete for overlapping binding sites on the kinesin heavy chain.

**Figure 2.**
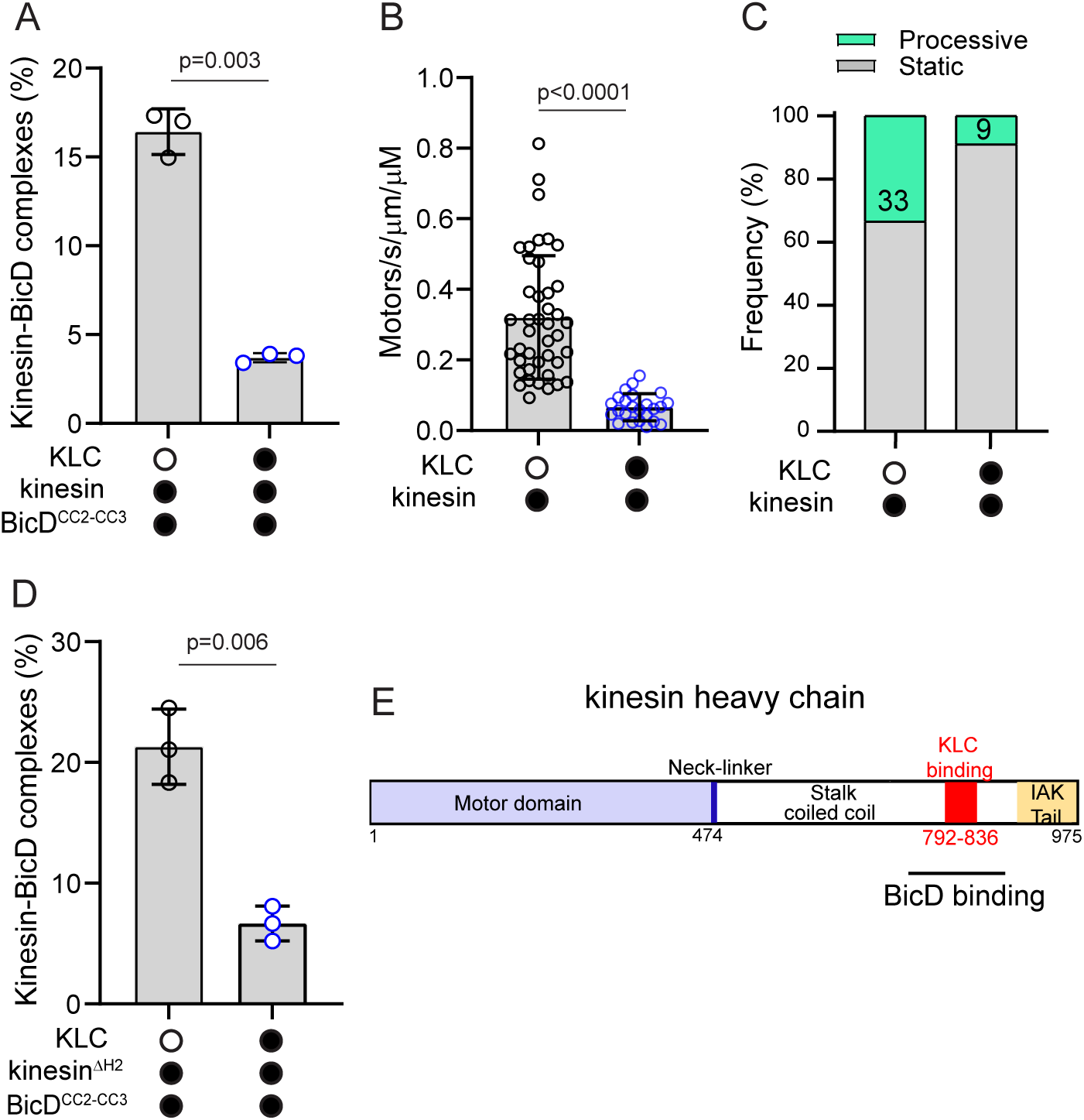
Kinesin light chain decreases the interaction of the kinesin heavy chain with BicD^CC2-CC3^. ***A***, Homodimeric kinesin without KLC forms more complexes with BicD^CC2-CC3^ than does heterotetrameric kinesin with bound light chains (KLC) (16.4 ± 1.3% versus 3.7± 0.26%, mean ±SD). Kinesin was separately expressed and purified with or without light chain. N=3 experiments. Unpaired t-test with Welch’s correction, p=0.003. ***B***, The presence of KLC decreased the number of kinesin motors that bound to the MT (0.32 ± 0.17 versus 0.07 ± 0.03, mean ± SD). N= 2 experiments. Number of MTs analyzed was n=41 without KLC and n=23 with KLC. Unpaired t-test with Welch’s correction, p<0.0001. ***C***, The presence of KLC reduced the fraction of motors moving processively from 33% to 9%. ***D***, The same experiment as in panel A, but using kinesin^ΔH2^ (21.3± 3.1% without KLC versus 6.6± 1.5% with KLC, mean ±SD), N=zz experiments. Unpaired t-test with Welch’s correction, p=0.006. A 5-fold molar excess of purified KLC was added to a preparation of kinesin^ΔH2^ expressed without KLC. ***E***, Schematic of the *Drosophila* kinesin heavy chain indicating that the region where the kinesin light chain (KLC) and BicD bind likely overlaps.

Alternatively, the weaker binding of BicD to kinesin^KLC^ could be an indirect result of the binding site for BicD being obscured in the auto-inhibited state. Heterotetrameric kinesin^KLC^ forms a more robust auto-inhibited state compared to kinesin heavy chain alone, as indicated by the heterotetramer showing fewer interactions with the MT than homodimeric kinesin (Fig. 2B) and fewer of those interactions resulted in processive runs (Fig. 2C). However, when we quantified the number of BicD-kinesin complexes obtained using kinesin^ΔH2^, which disfavors the inhibited state, we obtained very similar results to that seen with wild-type kinesin (compare Fig. 2*D* *versus 2A*). We conclude that the BicD and KLC binding sites on the kinesin heavy chain overlap (Fig. 2E), and that KLC plays a role in regulating the binding of kinesin heavy chain to BicD.

### BicD can bind two kinesins

When investigating the interaction between kinesin^ΔH2^ and BicD, we observed that the run length of kinesin ^ΔH2^ in the presence of BicD^CC2-CC3^ increased significantly from 1.8 ± 0.1 µm to 3.5 ± 0.1 µm (Fig. 3A). This finding raised the question of whether more than one kinesin can bind to BicD, because previous studies showed that the run length of two kinesins is longer than that of a single kinesin (36). To investigate the possibility that two kinesins bind per BicD, the biotin tag at the C-terminus of the kinesin^ΔH2^ heavy chain was labeled with either a 488 nm or a 647 nm Alexa dye, combined in equimolar ratios, and then added to BicD^CC2-CC3^ at a molar ratio of 2:1. We observed ∼7% dual-color dots, indicating that a small population (∼14%) of BicD molecules have two bound kinesins^ΔH2^. Single-colored images can be either kinesin^ΔH2^ alone or bound to BicD^CC2-CC3^. When the speed and run length of dual-colored complexes was compared to that of single-colored images, speed was unchanged (Fig. 3B), but the run length of dual-colored complexes was longer than that of single-colored images (Fig. 3C) showing functionally the effect of having two bound kinesins.

**Figure 3.**
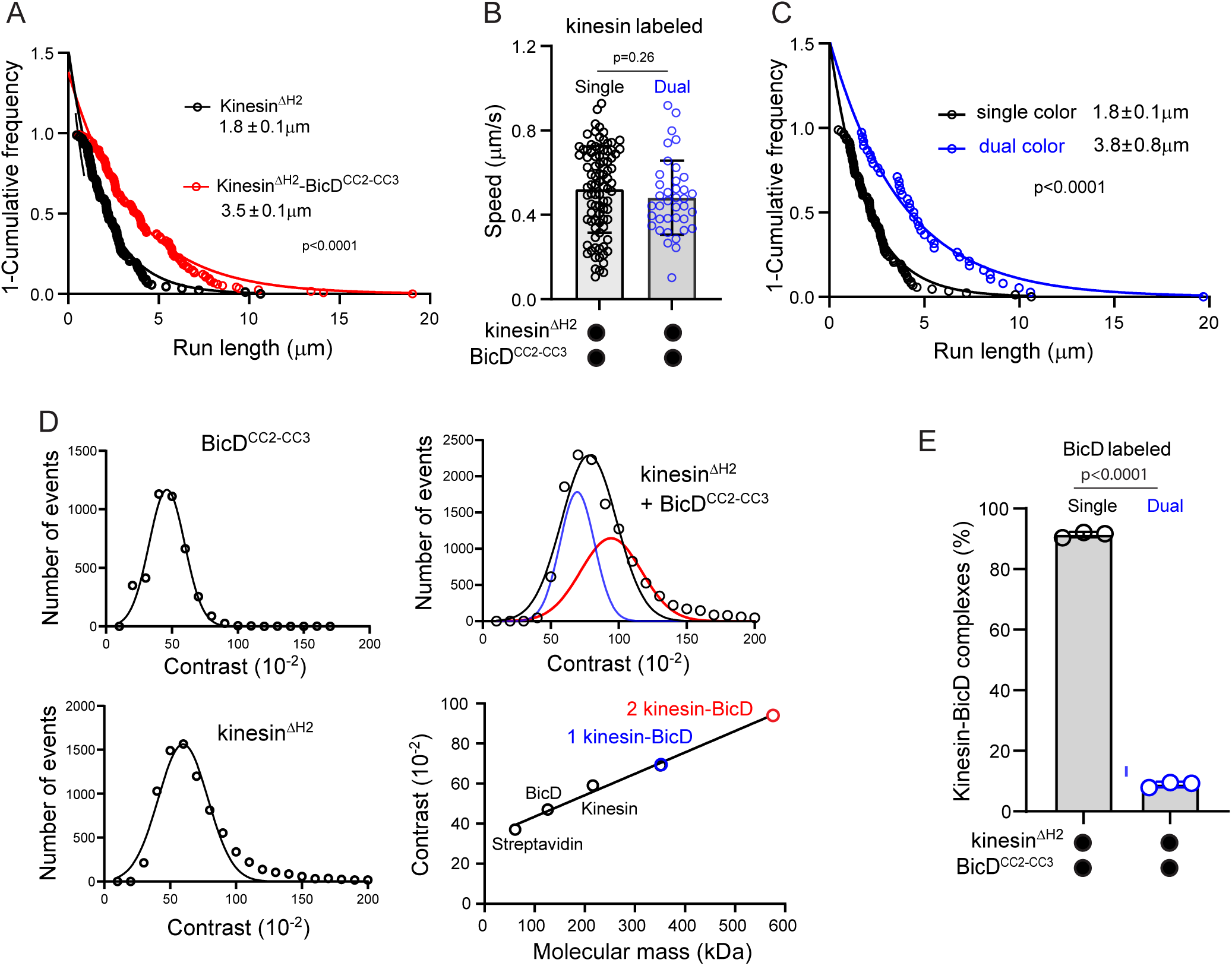
Two kinesins can bind to BicD. ***A*,** The run length of kinesin^ΔH2^ is enhanced from 1.8 ± 0.1 µm (n=88) to 3.5 ± 0.1 µm (n=115) in the presence of BicD^CC2-CC3^, N=3 experiments, Kolmogorov-Smirnov test, p<0.0001. ***B*,** Equimolar amounts of kinesin^ΔH2^ labeled with either Alexa 488 or Alexa 647 were mixed with BicD^CC2-CC3^. The speed of single-colored images (0.52 ± 0.20, n=88) was the same as that for dual-colored images with 2 kinesin^ΔH2^ bound (0.48 ± 0.17, n=38). N=2 experiments, unpaired t-test with Welch’s correction, p=0.26. ***C***, Run length of dual-colored images containing two kinesin^ΔH2^ complexes is longer (3.8 ± 0.8, n=38) than that of single-colored images (1.8±0.1, n=88, N=2 experiments, Kolmogorov-Smirnov test, p<0.0001). ***D***, Interferometric scattering mass spectrometry (iSCAMS) data showing contrast distributions for BicD^CC2-CC3^, kinesin^ΔH2^ and the kinesin ^ΔH2^-BicD^CC2-CC3^ complex, along with a standard curve of contrast as a function of molecular mass. BicD and kinesin were each fit to a single Gaussian and the kinesin-BicD complex histogram was fitted to the sum of two Gaussian distributions. ***E***, Equimolar amounts of BicD^CC2-CC3^ labeled with either 525 nm or 655 nm Qdots were mixed with kinesin^ΔH2^ and the number of red-green co-localizations quantified to show that a single kinesin does not bind to two BicD molecules. N=2 experiments, n=39 MTs. Unpaired t test with Welch’s correction, p<0.0001.

To more directly show that two kinesins can bind to BicD^CC2-CC3^, we measured the molecular mass of the kinesin^ΔH2^-BicD^CC2-CC3^ complex using interferometric scattering mass spectrometry (iSCAMS)(37) (Fig. 3D). A single Gaussian fitted the plot of number of events versus contrast for kinesin ^ΔH2^ alone (215 kDa with biotin tag) and for BicD^CC2-CC3^ alone (126 kDa with biotin tag). These two proteins, along with streptavidin were used to construct a standard curve of contrast versus molecular mass. The molecular mass of the kinesin ^ΔH2^-BicD^CC2-CC3^ complex is expected to be 340 kDa if one kinesin is bound and 556 kDa with two kinesins bound. The data fit a Gaussian distribution with a mean molecular mass of 430 kDa, which falls between the two possible outcomes. The data could also be fit to the sum of two Gaussian peaks, one centered at a mean molecular mass of 340 kDa (46% of the total area), and the other at 560 kDa (54% of the total area), suggesting that approximately half of the BicD^CC2-CC3^ molecules had two bound kinesins ^ΔH2^.

As a control we asked whether a single kinesin can bind to two BicD^CC2-CC3^ molecules. BicD^CC2-CC3^ was labeled with either 525 nm or 655 nm Qdots, mixed in equimolar ratios, and added to kinesin^ΔH2^. Very few dual-colored images were observed, showing that a single kinesin rarely binds to two BicD (Fig. 3E).

### BicD increases recruitment and the fraction of kinesin-1 motors moving processively on MTs

We next investigated the extent to which BicD activates kinesin from its auto-inhibited state. The motile properties (binding frequency to the MT, percent of bound motors that moved processively, speed and run length) of the kinesin-BicD^CC2-CC3^ complex were quantified relative to that of auto-inhibited kinesin or kinesin^ΔH2^. Auto-inhibited kinesin can bind to and move on MTs (Fig. 4A, left panel), but binding events were infrequent (Fig. 4B) and only 28% of these MT-bound events were processive (Fig. 4C). The speed of moving motors was slow (Fig. 4D) and run lengths were short (Fig. 4E). In contrast, the binding frequency for kinesin^ΔH2^ was significantly higher than for kinesin, and 48% of MT-bound motors moved processively (Fig. 4B-C). In addition, speed was 1.7-fold faster and run lengths 1.6-fold longer than for kinesin (Fig. 4D,E).

**Figure 4.**
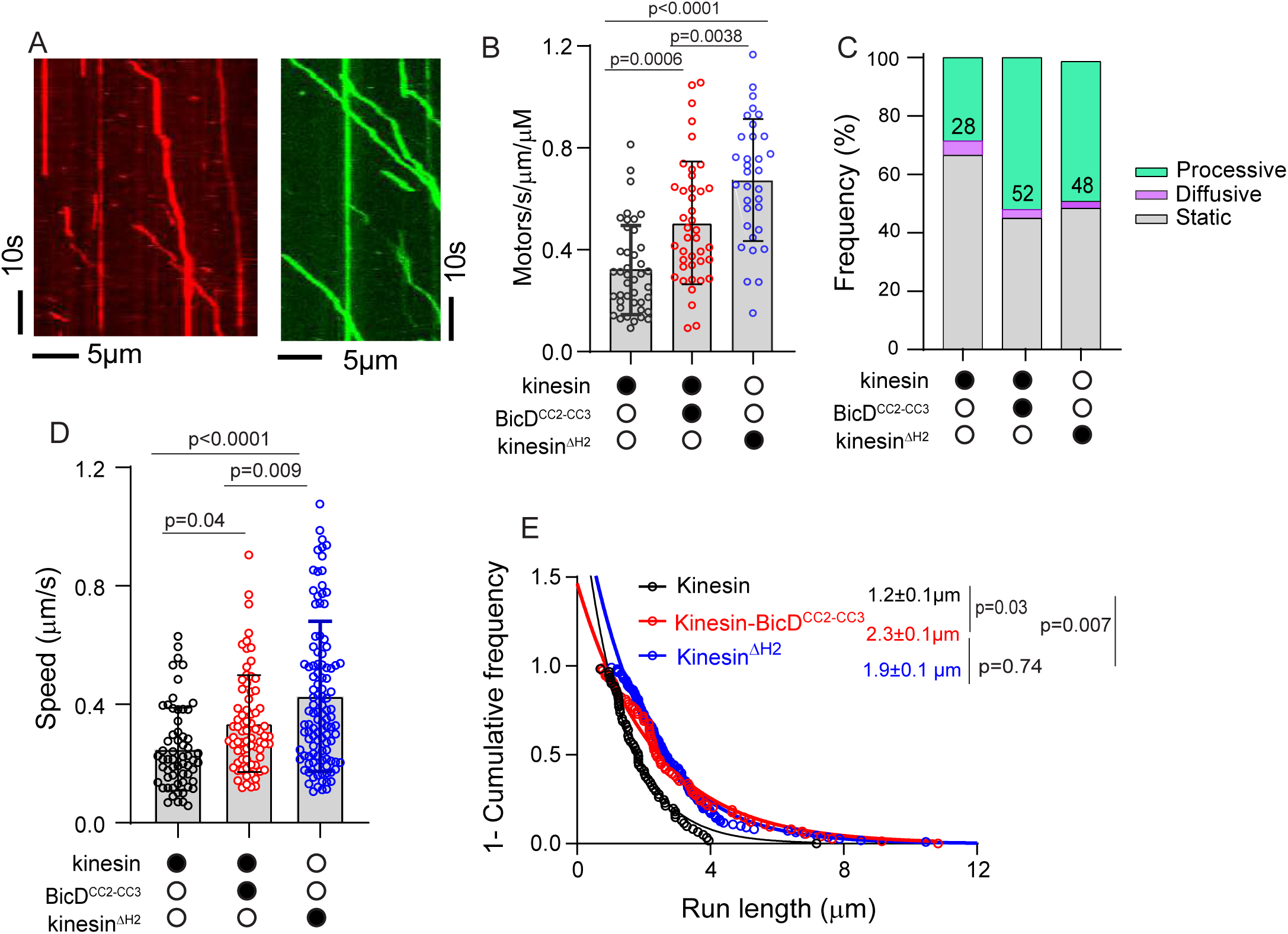
BicD increases the fraction of processively moving kinesin-1 motors. ***A***, Kymographs illustrating the trajectories of (*left*) kinesin or (*right*) the kinesin-BicD^CC2-CC3^ complex on MTs. ***B***, Recruitment of kinesin (motors/s/µm/µM) (0.32 ± 0.17, n=42 MTs), the kinesin-BicD^CC2-CC3^ complex (0.50 ± 24, n=41 MTs) and kinesin^ΔH2^(0.67 ±0.24, n=33 MTs) to the MT. Mean ± SD, N=3 experiments. One-way ANOVA with Tukey’s multiple comparison, p values indicated on figure. ***C***, Percentage of bound motors that move processively (green), diffusively (purple) or that were static (gray). The percentage of processive events were 28% for kinesin, 52% for the kinesin-BicD^CC2-CC3^ complex, and 48% for kinesin^ΔH2^. ***D***, Speed distributions of kinesin (0.25 ± 0.19 µm/s, n=74), the kinesin-BicD^CC2-CC3^ complex (0.33 ± 0.19, n=84) and kinesin^ΔH2^(0.43 ± 0.25 µm/s, n=111). Mean ± SD, N=3 experiments. One-way ANOVA with Tukey’s multiple comparison, p values indicated on figure. ***E***, Run length of kinesin (1.2 ± 0.1 µm, n=34), the kinesin-BicD^CC2-CC3^ complex (2.3 ± 0.1 µm, n=63) and kinesin^ΔH2^(1.9 ± 0.1 µm, n=79). N=3 experiments. One-way ANOVA with Tukey’s multiple comparison, p values indicated on figure.

For the kinesin-BicD^CC2-CC3^ complex, we first determined the optimal molar ratio of BicD^CC2-CC3^ to kinesin (Fig. S1). The highest percentage of processive events was observed at a 2:1 molar ratio, and thus a final ratio of 10 nM BicD:5nM kinesin was used for subsequent experiments. BicD^CC2-CC3^ was labeled with a Qdot, and kinesin was unlabeled. Compared to kinesin alone, the kinesin-BicD^CC2-CC3^ complex exhibited a 1.6-fold higher binding frequency to MTs, and the percentage of those events that were processive increased from 28% to 52%. Speeds were not significantly faster in the presence of BicD, but run lengths were increased 1.9-fold. Binding of kinesin to BicD^CC2-CC3^ thus activates motility relative to kinesin alone, consistent with BicD disrupting the auto-inhibited state of kinesin. The level of activation was similar to that observed with kinesin^ΔH2^ with respect to the percent of bound motors that moved processively and run length.

### MAP7 enhances kinesin recruitment to the MT with little change in processivity

We investigated the extent and mechanism by which the microtubule-associated protein *Drosophila* MAP7 affects kinesin function. MAP7 has two separable domains, an N-terminal region that binds MTs (MAP7^MTBD^) and a C-terminal region that binds kinesin (MAP7^KBD^) (Fig. 5A). Specifically, we determined if MAP7 enhances kinesin recruitment to the MT and/or the percent of bound kinesins that moved processively.

**Figure 5.**
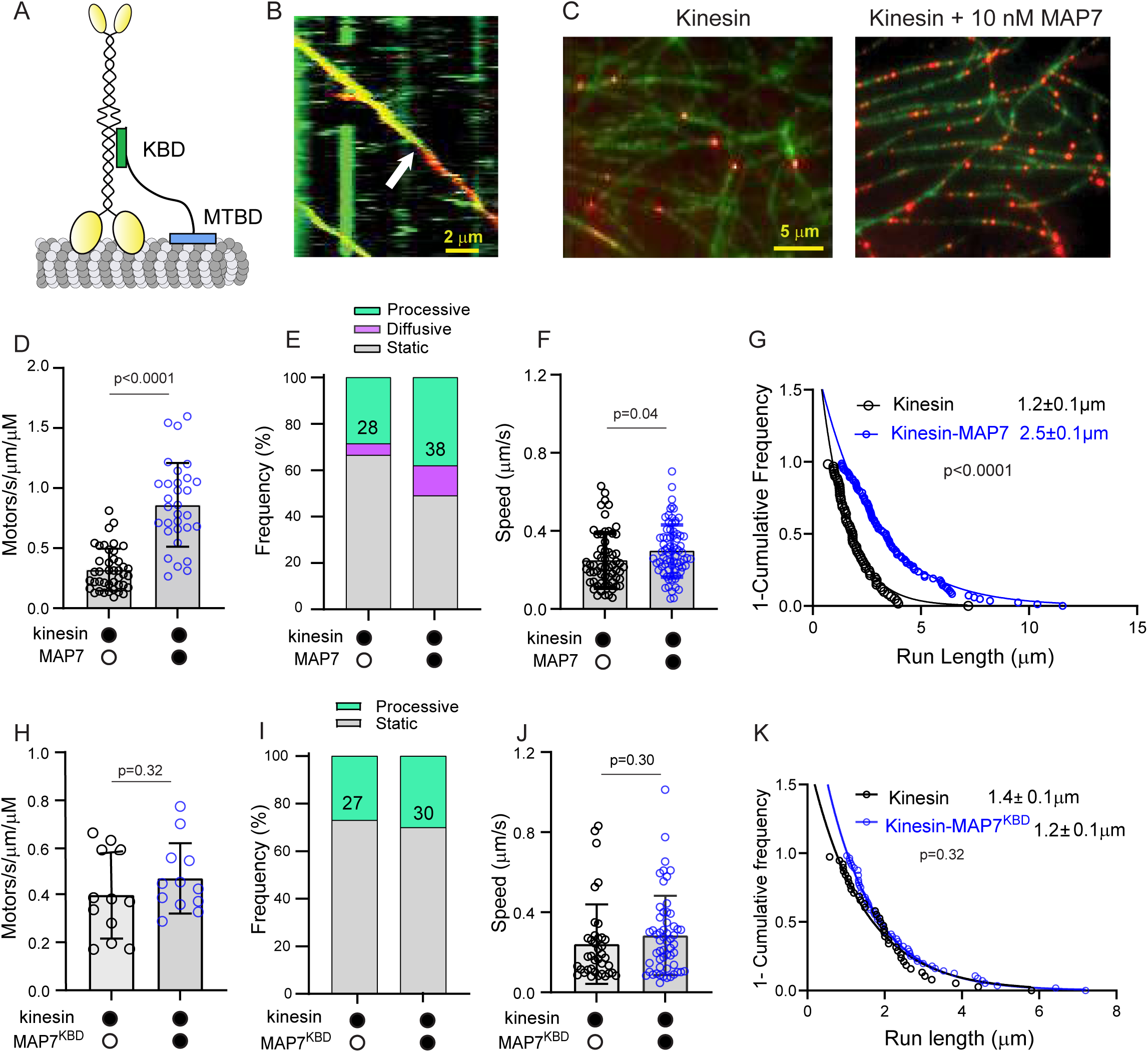
MAP7 enhances kinesin recruitment to the MT. ***A***, Schematic of the kinesin-MAP7 interaction. The kinesin-binding domain (MAP7^KBD^) and the microtubule-binding domain (MAP7^MTBD^) of MAP7 are indicated. ***B***, Kinesin was labeled with a red Qdot, and MAP7 with a green Qdot. The kinesin-MAP7 complex can move together processively on MT tracks (yellow), but MAP7 can also dissociate during the processive run (white arrow). ***C***, Fields illustrating enhanced recruitment of kinesin (red) to the MT (green) in the presence of 10 nM MAP7. ***D***, MAP7 enhances the number of kinesin motors bound to the MT from 0.32 ± 0.17 motors/s/µm/µM to 0.86 ± 0.35 (mean ± SD, N=2 experiments, n=42 and 31 MTs respectively. Unpaired t-test with Welch’s correction, p<0.0001. ***E***, The percentage of processive runs increases from 28% to 38% in the presence of MAP7, and the number of diffusive events increased from 5% to 13%. ***F***, The speed of kinesin in the presence of MAP7 (0.29 ± 0.13, n=76) is similar to that of kinesin alone (0.25 ± 0.14, n=61). Mean ± SD, N=2 experiments, Unpaired t-test with Welch’s correction, p=0.04. ***G***, The run length of the kinesin in the presence of MAP7 (2.5 ± 0.1 µm) is significantly longer than that of kinesin alone (1.2 ± 0.1µm). N=2 experiments, Kolmogorov-Smirnov test, p<0.0001. ***H***, MAP7^KBD^ did not enhance the number of kinesin motors bound to the MT (0.40 ± 0.18 motors/s/µm/µM versus 0.47 ± 0.15, n=12 MTs, mean ± SD, N=2 experiments Unpaired t-test with Welch’s correction, p=0.32). ***I***, The percentage of processive runs was unchanged in the presence of MAP7 ^KBD^ (27% versus 30%). ***J***, The speed of kinesin in the presence of MAP7^KBD^ (0.29 ± 0.13, n=76) is not significantly different from that of kinesin alone (0.25 ± 0.19, n=74). Mean ± SD, N=2 experiments, Unpaired t-test with Welch’s correction, p=0.30. K, The run length of the kinesin-MAP7^KBD^ complex (1.2 ± 0.1 µm) was not significantly different than that of kinesin alone (1.4 ± 0.1µm). N=2 experiments, Kolmogorov-Smirnov test, p=0.32.

To first establish that the two proteins interact, kinesin and MAP7 were labeled with different color Qdots. The complex could occasionally move processively together on MTs, but MAP7 could also detach from kinesin during a processive run (Fig. 5B), suggesting that the interaction between these two proteins is dynamic. The optimal MAP7 concentration to use with 5 nM kinesin was determined to be 10 nM based on the quantification of the percentage of bound kinesins that moved processively (Fig. S2, A-H).

The two parameters that were most affected by MAP7 were the number of kinesins recruited to the MT and the run length. MAP7 (10 nM) enhanced the number of kinesin motors bound to the MT by 2.7-fold (Fig. 5C,D), and the run length of the kinesin-MAP7 complex was 2.1-fold longer than that of kinesin alone (Fig. 5G), consistent with MAP7 acting as a tether to recruit and then prevent kinesin dissociation from the MT. The percentage of processive runs, however, only increased from 28% to 38% in the presence of MAP7, accompanied by a larger percentage of diffusive events (13%) (Fig. 5E) than seen with kinesin or kinesin^ΔH2^ (see Fig. 4C). Kinesin’s speed was only marginally changed by MAP7 (Fig. 5F).

We further investigated whether the C-terminal half of MAP7 (MAP7^KBD^) affected any of the motility parameters observed with the full-length molecule. Given that the domain that directly binds to kinesin is the one that could most likely impact the auto-inhibited state of kinesin, we were most interested in the percentage of motors that moved processively. At 10 nM, the same concentration used with full-length MAP7, there was no enhanced recruitment to the MT, no increase in the percent of motors that moved processively, and no enhancement in speed or run length (Fig.5, H-K). There was no change in the number of kinesins bound to the MT nor in the percent of motors that moved processively over a range of MAP7^KBD^ concentration from 1-50 nM, and only a small enhancement in both parameters at 100 nM, 10-times the concentration used with full-length MAP7 (Fig. S2, I-J).

Taken together, these results demonstrate that the two domains of MAP7, MAP7^MTBD^ and MAP7^KBD^, act in concert to enhance kinesin recruitment to the MT and prolong run lengths, while playing a smaller role in releasing kinesin from its auto-inhibited state.

### Kinesin motility is most active in the presence of both MAP7 and BicD

Lastly, we investigated whether the combination of BicD and MAP7 together would have the largest effect on kinesin motility. MAP7, BicD^CC2-CC3^ and kinesin were mixed at a molar ratio of 2:2:1. We analyzed recruitment to the MT, percentage of processive motion, speed and run length and compared it with values measured previously in this study. The binding frequency of kinesin to MTS in the presence of both MAP7and BicD^CC2-CC3^ was the same as with MAP7 alone, but higher than for kinesin, kinesin^ΔH2^, or the kinesin-BicD^CC2-CC3^ complex (Fig. 6A). Both the percent processive motion and run length were higher than seen with either BicD^CC2-CC3^ or MAP7 alone (Fig. 6B, D). Speed was relatively unchanged for all combinations except for the lower speed seen with auto-inhibited kinesin (Fig. 6C). Taken together, our findings are consistent with the notion that interaction with multiple binding partners allows kinesin to function as the most efficient cargo transporter.

**Figure 6.**
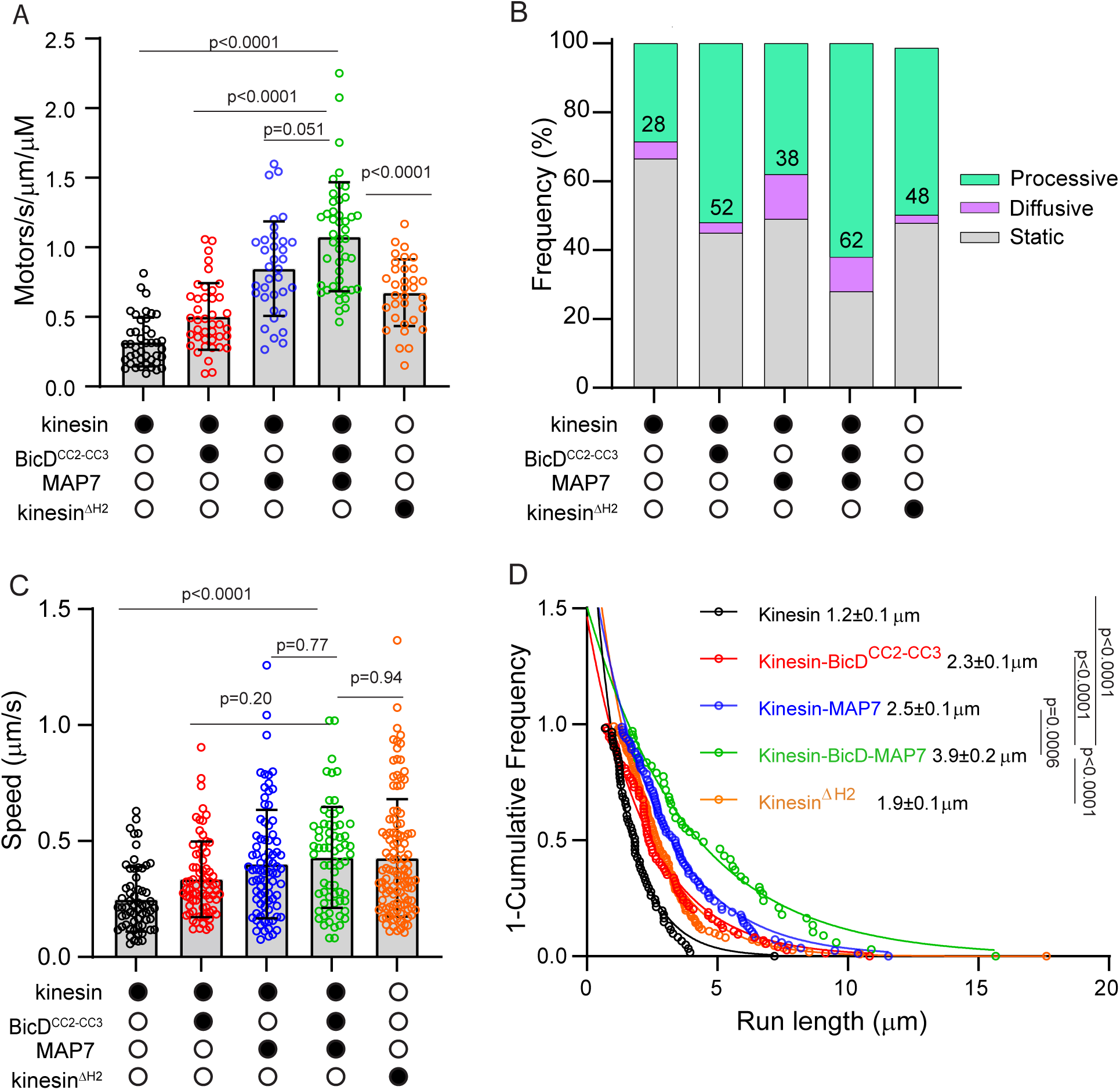
The combination of BicD and MAP7 produces the most active kinesin. Values for the kinesin-BicD^CC2-CC3^-MAP7 complex are shown and compared to data previously shown with other kinesin or kinesin-complexes. ***A***, The combination of MAP7 and BicD^CC2-CC3^ enhances the number of kinesin motors bound to microtubules (from 0.32 ± 0.17 motors/s/µm/µM to 1.1 ± 0.39, N=3 experiments, n=45 MTs, One-way ANOVA with Tukey’s multiple comparison. p values indicated on figure). ***B***, The percentage of processive runs was 62% in the presence of both MAP7 and BicD^CC2-CC3^, higher than with either binding partner alone. ***C***, The speed of kinesin in the presence of both MAP7 and BicD^CC2-CC3^ is 0.43±22 µm/s, n=67. N=3 experiments, One-way ANOVA with Tukey’s multiple comparisons are shown with p values, ***D***, The run length of kinesin in the presence of both MAP7 and BicD^CC2-CC3^ (3.9 ± 0.2 µm) was statistically longer than all other variations. N=3 experiments, One-way ANOVA with Tukey’s multiple comparison.

## Discussion

Here we investigated the extent to which BicD and MAP7 activated auto-inhibited kinesin-1, separately and in combination. The two parameters that were most revealing were the number of kinesins recruited to the MT (per time per MT length), and the percent of those bound motors that exhibited processive motion. We interpret an increase in the percent of processive motors to reflect abolishment of the auto-inhibited state of kinesin. An increase in the number of motors bound to the MT can be either a result of motor domains being more available for binding to the MT when kinesin is not in the auto-inhibited state, or due to an independent recruitment mechanism. We show that auto-inhibited homodimeric *Drosophila* kinesin is activated by binding to *Drosophila* BicD, resulting in more kinesins binding to the MT and more of those bound motors exhibiting processive motion. *Drosophila* MAP7, in contrast, predominantly recruits more kinesins to the MT and enhances the run length of those motors. Similar results were obtained in an earlier study which showed that MAP7 increased the number of processive and diffusive events to a similar degree, consistent with recruitment to the MT but not relief of auto-inhibition (34). In combination, these two binding partners cause kinesin to be similar in activity to a constitutively active full-length kinesin lacking hinge 2, with the additional benefit of a longer run length courtesy of MAP7.

Enhanced recruitment to the MT and longer run lengths were observed at 10 nM MAP7. The effect of MAP7 on motility is highly concentration-dependent because the binding site for MAP7 on the MT partially overlaps with the kinesin binding site (Ferro Yildiz Science 2022). In contrast, MAP7^KBD^ (the C-terminal region of MAP7 that binds the kinesin stalk but does not interact with MTs), had no significant effect on either parameter at 10 nM. A weak binding to kinesin is necessary for MAP7 to recruit the motor to the MT, but not act as an anchor to prevent subsequent motion. At 10-fold higher concentration (100 nM), MAP7^KBD^ did show an enhancement in the percent of processive motors, but at this concentration the full-length MAP7 would be inhibitory. A recent study (35) also reported a higher run frequency with 1-5 µM of the kinesin binding domain of MAP7, while the full-length MAP7 showed positive effects in the range of 10 nM and negative effects at higher concentrations due to competition for MT binding sites. Only if the “effective concentration” of the MAP7^KBD^ becomes higher when both kinesin and MAP are bound to the MT lattice would our data support the idea that MAP7 plays a large role in relieving auto-inhibition of kinesin. The auto-inhibited state of kinesin is stabilized by multiple weak interactions along the stalk (27, 28). Given that MAP7^KBD^ binds to the coiled-coil stalk adjacent to the region where hinge 2 and the kinesin heavy chain-light chain interface abut in the inhibited state, it would not be surprising if MAP7 binding could contribute to weakening the auto-inhibited state.

The essential nature of MAP7 was highlighted by studies in *Drosophila*, which showed that flies lacking MAP7 do not survive until adulthood, and that peroxisome motion in cultured *Drosophila* neurons was severely impaired in either kinesin null or MAP7 null cells (30). In addition, the kinesin binding domain of MAP7 was sufficient to sustain mitochondrial motility in S2 cells depleted of endogenous MAP7, indirectly suggesting that it may play a role in overcoming auto-inhibition of kinesin (30). More recent studies, however, suggest that biological context may influence whether the kinesin binding domain of MAP7 is sufficient to affect kinesin function. While expression of the kinesin binding domain alone rescues centrosome separation defects in *Drosophila* brain neuroblasts, it did not restore mRNA localization or fast oocyte streaming (32).

Our results and interpretation align with a recent study by McKenney and colleagues (29), which suggest that human hetero-tetrameric kinesin is activated by synergistic mechanisms. In their study, a combination of binding of the cargo-adaptor protein nesprin-4 to the light chain of kinesin-1 in combination with MAP7 result in the most active kinesin. A notable difference in our study is that BicD binds to the heavy chain of homodimeric kinesin for relief of auto-inhibition, and kinesin light chains decrease binding to BicD. KLC can thus be considered a modulator of kinesin function, as it dictates the degree of binding to BicD and thus indirectly how activated the pool of available kinesin will be. In both studies, MAP7 was shown to recruit more motors to the MT, without directly having an effect on release of auto-inhibition of kinesin. In a broad sense, both release of auto-inhibition and recruitment of more motors to the MT can be considered as “activation”, but here we make a distinction between the two mechanisms. A multi-step activation process may allow finer tuning of activity levels than one process alone.

We also show that BicD has the capacity to bind two kinesins, which is a further way to obtain a larger spectrum of activities that can modulate cargo movement to the plus end of the MT. Longer run lengths were seen when two kinesins were bound to BicD, which presumably would also result in higher force production. The ability of BicD to bind two kinesins mirrors our previous study that demonstrated recruitment of two dyneins to BicD (13), similar to observations by others using mammalian BicD2 (17). BicD with two bound dyneins moved faster and longer than BicD with only one dynein. Thus, a consequence of two kinesins being able to bind to BicD would be maintenance of more balanced forces between kinesin and dynein during cargo transport.

Structural evidence is needed to reveal whether kinesin binds to two different regions in the central domain of BicD, or whether each chain of the coiled-coil can bind a kinesin via the same primary sequence. Based on the AlphaFold protein structure database, the kinesin binding site is located in helices 3 and/or 4 of BicD (shown schematically in Fig. 1A). The binding of mammalian kinesin to BicD2 was also mapped to the central region of BicD2 (38). Thus both the Drosophila and mammalian BicD family proteins have three domains: an N-terminal region that binds dynein-dynactin, a central region that binds kinesin, and a C-terminal region that binds cargo-adaptor proteins to link the motor complex to cargo.

A number of dynein activating adaptors bind both dynein-dynactin and kinesin. Although dynein-dynactin is uniformly activated by these adaptors, their effect on kinesin activity has not been investigated in as much detail. A recent study using cellular lysates showed that a full-length activating adaptor TRAK2, which links both dynein and kinesin-1 motors to mitochondria, strongly activates kinesin-1 (39). In contrast, purified truncated constructs of TRAK1 and TRAK2 were insufficient to robustly activate homodimeric kinesin-1 unless MAP7 was also present (40). These disparate results require further investigation to be reconciled. Another example of kinesin binding to a dynein activating adaptor is Hook3 and kinesin-3 (KIF1C). Kinesin-3 isolated from human 293T cells was a robust processive motor in the absence of Hook3, and addition of Hook3 resulted in no further activation (41). In contrast, Hook3 doubled the MT recruitment of kinesin-3 that was expressed in insect cells, suggesting a release of auto-inhibition upon binding to Hook3 (42). The reasons for this discrepancy has not been resolved, but could be due to differences in post-translational modifications to the motor or other factors. It is not a universal feature that all activating adaptors allow simultaneous binding of dynein and kinesin; for example, an active dynein-dynactin TRAK1 complex that moves toward the minus end of the MT was reported to be incompatible with kinesin binding (40).

This study lays the groundwork for investigating the motion of biologically relevant complexes containing both dynein and kinesin-1 linked via BicD. The goal is to understand what factors bias the direction of motion of the complex, and whether bidirectional motion caused by the active motor switching from dynein to kinesin will be observed during a trajectory. It was hypothesized based on studies with isolated phagosomes and mathematical modeling that the net transport of cargoes can be regulated by controlling only one motor, in particular kinesin-1 (34). Knowing that one or two kinesins can bind to BicD, and that MAP7 enhances kinesin recruitment to the MT, will allow us to test directional outcomes in a systematic way.

## Experimental procedures

### Kinesin-1 expression and purification

The *Drosophila melanogaster* full-length kinesin heavy chain (accession number AF053733) was expressed in Sf9 cells with a C-terminal biotin tag, which is an 88-amino acid segment derived from the E. coli carboxyl carrier protein, followed by a FLAG tag to facilitate purification. Biotinylation occurs during expression in Sf9 cells (43). The kinesin light chain KLC2 (accession number AF055298) was cloned with a 6XHis tag at its C-terminus. Recombinant baculovirus was prepared by standard protocols.

Kinesin heavy chains and light chains were co-expressed in Sf9 cells and grown for approximately 72 hours at 27°C in media supplemented with 0.2 mg/ml biotin, harvested by centrifugation and re-suspended in 10 mM imidazole, pH 7.0, 0.3 M NaCl, 1 mM EGTA, 5 mM MgCl_2_, 7% sucrose, 2 mM DTT, 0.5 mM AEBSF, 0.5 mM TLCK, 5 μg/ml leupeptin, and 1 mM ATP. Cells were lysed by sonication, and the lysate was clarified by centrifugation at 200,000 x g for 30 min. The supernatant was applied to a FLAG affinity resin column (Sigma-Aldrich, A2220) and washed with 10 mM imidazole, pH 7.0, 0.3 M NaCl, and 1 mM EGTA. Kinesin was eluted from the column by addition of 0.1 mg/ml FLAG peptide (APEXBIO, A6002) to the buffer. Eluted fractions of interest were supplemented with 1 mM DTT, 1 μg/ml leupeptin, and 10 μM ATP before being concentrated using an Amicon centrifugal filter device (Millipore, UFC801024). Kinesin was clarified at 487,000 x g for 20 min and dialyzed versus 10 mM imidazole, pH 7.4, 200 mM NaCl, 55% glycerol, 1 mM DTT, 10 µM MgATP, and 1 µg/ml leupeptin. Protein concentration was determined with the Bradford reagent (Thermo Scientific, 1856210), using BSA as a standard. The protein was flash frozen in small aliquots and stored at −80°C. *Drosophila* kinesin^ΔH2^ (lacking amino acids K521-D641) with a C-terminal biotin and FLAG tag was similarly expressed and purified.

### BicD expression and purification

Full-length *Drosophila* BicD, BicD^CC1^(amino acids 21-380), BicD^CC2^(amino acids 318-557), BicD^CC3^(amino acids 536-782), BicD^CC2-CC3^ (amino acids 318-782), BicD^318-658^, BicD^437-782^ were expressed in Sf9 cells for approximately 72 hours at 27°C in media supplemented with 0.2 mg/ml biotin (Sladewski et al. 2018). Each construct had a FLAG tag followed by a biotin tag at the N-terminus. Purification was the same as described above for kinesin, except that ATP was omitted.

### MAP7 expression and purification

*Drosophila melanogaster* MAP7 (accession number NP_728941.2), with N-terminal FLAG and SNAP tags, was expressed in Sf9 cells as described above for BicD. MAP7^KBD^ (amino acids 499-887) was expressed in BL21(DE3) bacterial cells. Cells were induced with 1.0 mM isopropyl 1-thio-D-galactopyranoside and grown at 37 °C for 3 hr. Cells were pelleted by centrifugation and resuspended in 10 mM sodium phosphate, pH 7.5, 0.3 M NaCl, 0.5% glycerol, 7% sucrose, 7 mM-mercaptoethanol, 5 µg/ml leupeptin, 10 mM AEBSF, 0.5 mM phenylmethylsulfonyl fluoride, and 0.5 mM TLCK. The lysate was sonicated, clarified by centrifugation, and the supernatant applied to a HIS-Select nickel affinity column (Sigma-Aldrich P6611). The column was washed with 10 mM sodium phosphate, pH 7.5, 0.3 M NaCl, 10 mM imidazole, and then the same buffer containing 30 mM imidazole. The protein was eluted with 10 mM sodium phosphate, pH 7.5, 0.3 M NaCl, and 0.2 M imidazole. The fractions of interest were combined and concentrated by dialysis in buffer containing 10 mM HEPES, pH 7.5, 0.3 M NaCl, 1 mM DTT, and 1µg/ml leupeptin and 50% glycerol. Protein concentration was determined with the Bradford reagent (Thermo Scientific, 1856210), using BSA as a standard. Protein was stored at −20 °C.

### Rigor Kinesin expression and purification

Rigor kinesin (G235A mutation in rat kinesin 1-406, accession number XP_032759158) with an N-terminal 6xHIS tag was cloned into pET24b and expressed in E. coli Rosetta (DE3) (Millipore-Sigma 70954). Cells were induced with 1.0 mM isopropyl 1-thio-D-galactopyranoside and grown at 25 °C for 18 hr. Cells were pelleted by centrifugation and resuspended in lysis buffer (10 mM sodium phosphate, pH 7.5, 0.2 M sodium chloride, 1 mM ethylene glycol-bis(2-aminoethylether)-NNN’N’-tetraacetic acid, 0.5 mM dithiothreitol, 0.5 mM AEBSF, 5 µg/ml leupeptin, 0.5 mM TLCK, and 0.5 mM PMSF). The lysate was sonicated, clarified by centrifugation and applied to a HIS-Select nickel affinity resin (Sigma-Aldrich, P6611). The resin was washed with 10 mM sodium phosphate, pH 7.5, 0.2 M sodium chloride, 1 mM ethylene glycol-bis(2-aminoethylether)-NNN’N’-tetraacetic acid, 0.5 mM dithiothreitol containing 10 mM imidazole, then containing 30 mM imidazole, and protein eluted with the same buffer containing 0.2 M imidazole. The protein was pooled and concentrated using an Amicon centrifugal filter device (Millipore, UFC801024), and dialyzed versus 10 mM imidazole, pH 7.5, ).2 M sodium chloride, 1 mM ethylene glycol-bis(2-aminoethylether)-NNN’N’-tetraacetic acid, 50% glycerol, 0.5 mM DTT and 1 µg/ml leupeptin. Protein concentration was determined with the Bradford reagent (Thermo Scientific, 1856210), using BSA as a standard. Protein was stored at −20°C.

### Microtubules

To prepare labeled microtubules for the motility assay, unlabeled tubulin isolated from cow brains was mixed with rhodamine-labeled tubulin (Cytoskeleton, Denver, CO, TL590M) at a 1:10 molar ratio on ice. The tubulin mixture was polymerized by incubation in a 37°C water bath for 30 min and then stabilized with 10 mM paclitaxel (Cytoskeleton, Denver, CO TXD01). Stabilized microtubules were kept at room temperature.

### Flow cell preparation

Glass cover slides measuring 24×60 mm (Fisher Scientific, 12541037) were plasma cleaned for 5 min on each side and were subsequently placed in glass Coplin jars. These jars were filled with 1M KOH and immersed in a sonicating water bath for 30 minutes. Following this, the slides were washed with water and then 95% ethanol. Once dried, the slides were moved to Coplin jars containing a solution of 1.73% 2-methoxy(polyethyleneoxy)propyltrimethoxysilane (Gelest, Inc., SIM6492) and 0.62% n-butylamine (Acros Organics, 109-73-9) in anhydrous toluene (Sigma-Aldrich, 244511). The slides were incubated for 90-120 min at room temperature. After incubation, the slides were sequentially immersed in two beakers filled with anhydrous toluene to remove excess chemicals. The slides were then dried using a nitrogen stream and stored at 4°C.

To ensure that the microtubules would not depolymerize, all buffers were equilibrated at room temperature. The PEGylated glass slides were coated with 0.2 mg/ml rigor kinesin for microtubule attachment, and then rinsed 2-3 times with BRB80 buffer containing 80 PIPES (Sigma-Aldrich P6757), pH 7, 1 mM MgCl_2_ (Quality Biological Inc. 351-033-721) 1 mM EGTA (Sigma-Aldrich, E4378) to remove excess rigor kinesin. Rhodamine-labeled microtubules were then introduced to the flow cell, and allowed to attach to the glass surface via the rigor kinesin for 5 min. The flow chamber was subsequently washed 2-3 times with BRB80 containing 10 mM DTT (Fisher Scientific ICN10059750), 5 mg/ml BSA (Sigma-Aldrich A3069), 0.5 mg/ml κ-casein (Sigma-Aldrich P-C0406) and 0.5% Pluronic F68 (Sigma-Aldrich P-1300) to remove any excess MTs and to block the glass surface with BSA and casein. Finally, the samples were diluted in motility buffer (80 PIPES (Sigma-Aldrich P6757), pH 7, 1 mM MgCl_2_, 1 mM EGTA, 10 mM DTT (Fisher Scientific ICN10059750), 5 mg/ml BSA (Sigma-Aldrich, A3069), 0.5 mg/ml κ-casein (Sigma-Aldrich, P-C0406) and 0.5% Pluronic F68 (Sigma-Aldrich, P-1300), and 2 mM MgATP (Sigma-Aldrich, A2383)) and observed for motion using TIRF microscopy.

### Formation of the Kinesin-BicD complex

Kinesin and BicD constructs were diluted into BRB80 buffer (80 mM PIPES, pH 7.0, 2 mM MgCl_2_, 0.5 mM EGTA) with freshly added 20 mM DTT and clarified for 20 min at 400,000 × g. Protein concentration was determined with the Bradford reagent (Thermo Scientific, 1856210) using BSA as a standard. In separate tubes, kinesin was mixed with 525 nm streptavidin Qdots (Invitrogen, Q10141MP) in a 1:2 molar ratio and BicD was mixed with 655 nm streptavidin Qdots (Invitrogen, Q10121MP) in a 1:1 molar ratio in separate tubes, and the mixture was incubated on ice for 15 min. To block any excess binding sites on streptavidin Qdots, 5 µM biotin was added to both tubes. The labeled kinesin and BicD were then mixed together in a molar ratio of 1:2 (500 nM kinesin, 1 µM BicD) and incubated on ice for 30 min in BRB80, 20 mM DTT (Fisher Scientific, ICN10059750), 2 mM MgATP (Sigma-Aldrich, A2383), 2.5 mg/ml BSA (Sigma-Aldrich, A3069), 0.5 mg/ml kappa-casein (Sigma-Aldrich, P-C0406), 0.5% Pluronic F68 (Sigma-Aldrich, P-1300), 10 µM paclitaxel (Cytoskeleton, Denver, CO TXD01), and an oxygen scavenger system (5.8 mg/ml glucose (EM Science, DX0145), 0.045 mg/ml catalase (Sigma-Aldrich, C40), and 0.067 mg/ml glucose oxidase (Sigma-Aldrich, G6125). The kinesin-BicD complex was diluted into motility buffer, and the movement of kinesin on microtubules was observed at a final kinesin concentration of 5 nM. To further confirm the interaction between kinesin and various BicD constructs, BicD was labeled with streptavidin Qdots in 1:1 molar ratio while kinesin was kept unlabeled, and the motion on microtubules was observed by TIRF microscopy.

### Determination of the number of kinesin molecules bound to BicD

Kinesin^ΔH2^ (2μl of 2 μM) were separately labeled with either 488 nm, or 647nm Alexa fluor (2 μl of 2 μM) in buffer BRB80 and incubated for 15 min on ice. The labeled kinesins with mixed in a 2:1 molar ratio with unlabeled BicD^CC2-CC3^ and incubated for 30 min on ice. The two-motor complex was diluted in the motility buffer described above and observed on a glass surface using TIRF microscopy. For motility assays, kinesin^ΔH2^ was separately labeled with either 525 nm or 655 nm streptavidin Qdots and then mixed with BicD. The motion of the two-motor complex was observed by TIRF microscopy.

Interferometric scattering microscopy was used to determine the molecular mass of the kinesin-BicD complex, using the procedure outlined in (37). Specifically, kinesin^ΔH2^ and BicD^CC2-CC3^ were combined in a 2:1 molar ratio in filtered BRB80 buffer. Streptavidin, BicD and kinesin were used to create a standard curve for contrast/mass conversion. Protein was diluted to 1 nM and flowed into a clean flow cell and mounted onto a Refeyn One^MP^ interferometric scattering microscopy (iSCAT) (Refeyn Ltd. Unit 9, Trade City, Sandy Ln W, Oxford OX4 6FF, UK) for observation. Movies of events of protein binding to the surface were recorded for 1 min and later exported for processing by software to find the contrast of individual binding events and converted to mass using the standard curve. The histogram of mass of proteins that binds to the surface was fit to Gaussian distribution to find the molecular mass of protein complexes.

### Formation of the kinesin-MAP7 complex

Kinesin was labeled with 565 nm streptavidin Qdots and then mixed with varying concentrations of MAP7(5-80 nM). After 30 min incubation on ice, the samples were observed on MTs using TIRF microscopy. To directly observe the interaction between kinesin and MAP7, each protein was labeled with a different color Qdot (525 nm or 655 nm). Two μl of 1 μM kinesin was mixed with 4 μl of 1 μM Qdot, and 2 μl of 2 μM MAP7 was mixed with 2 μl of 1 μM Qdot. After 15 min incubation, 5 μM biotin was added to each tube to block unoccupied binding sites on the streptavidin Qdots, followed by an additional 15 min incubation. Subsequently, the Qdot-labeled kinesin and MAP7 were mixed and incubated for another 30 min. Following dilution to 5 nM kinesin, the motion on MTs was observed by TIRF microscopy.

### Microscopy and data analysis

The single molecule images were acquired on a Nikon ECLIPSE Ti microscope equipped with objective-type TIRF. Qdots (525 nm or 655 nm) were excited using a 488 nm laser line, while the Alexa fluor dyes were illuminated by 488 nm or 647 nm laser lines. Additionally, a 561 nm laser line was used to illuminate rhodamine-labeled microtubules. Two Andor EMCCD cameras (Andor Technology USA, South Windsor, CT) were employed to capture 300-600 frames at 200 ms intervals (5 frames/s) for moving complexes. All experiments were conducted at a room temperature of 23 ± 1°C. To detect the immobile dual color complex on the glass surfaces, 30-60 frames at 200 ms intervals were captured. The resolution of the captured images was 0.1066 µm/pixel. The MTrackJ plugin of Image J was utilized to track moving objects in image sequences and provide data for further analysis. The travel distances or run lengths were measured from the appearance of the Qdot-labeled kinesin until its detachment from the microtubules. Run length data were plotted as a 1-cumulative probability distribution. GraphPad Prism Software (Version 10) was used to fit the data to a one-phase exponential decay equation p(x) = Ae-x/λ, where x denotes the travel distance along microtubules, p(x) represents the relative frequency, and A represents the amplitude. The speed was calculated by dividing the total length by the total time and reported as mean ± SD, while the run length was reported as mean ± standard error (SE). To calculate processive events or run frequency, the total number of runs per s per microtubule length was counted. To determine the statistical significance between two sets of run-length data, the nonparametric Kolmogorov-Smirnov test was conducted with 95% confidence interval using GraphPad Prism software (Version 10). An unpaired t-test was performed to calculate the *p*-value between two sets of speed data. A *p*-value equal to or below 0.05 is considered to be statistically significant. For three or more run length or speed or run frequency datasets, statistical significance was calculated using one-way ANOVA followed by Tukey’s post-hoc test.

### Criteria for processive versus diffusive motion

If kinesin traveled a distance greater than 0.5 µm, occasionally exhibiting diffusive motion but with a bias towards the plus end of the microtubule, it was classified as processive motion. When kinesin traveled distances up to 0.5 µm and displayed a back-and-forth motion resulting in an approximate zero displacement, it was categorized as diffusive motion.

## Data availability

All data are included in the article or supporting information. Raw data for each figure are provided as an Excel spreadsheet in Supporting Information

## Supporting Information

This article contains supporting information.

## Acknowledgements

We thank Elena Krementsova and Carol Bookwalter for their contributions to cloning and protein expression and purification.

## Funding and additional information

This work was supported by NIH grant R35 GM136288 to KMT and NIH grant R03 NS126811-01A1 to MYA.

## Conflict of interest

The authors declare that they have no conflicts of interest with the contents of this article.

---

**Supporting Figure 1.**
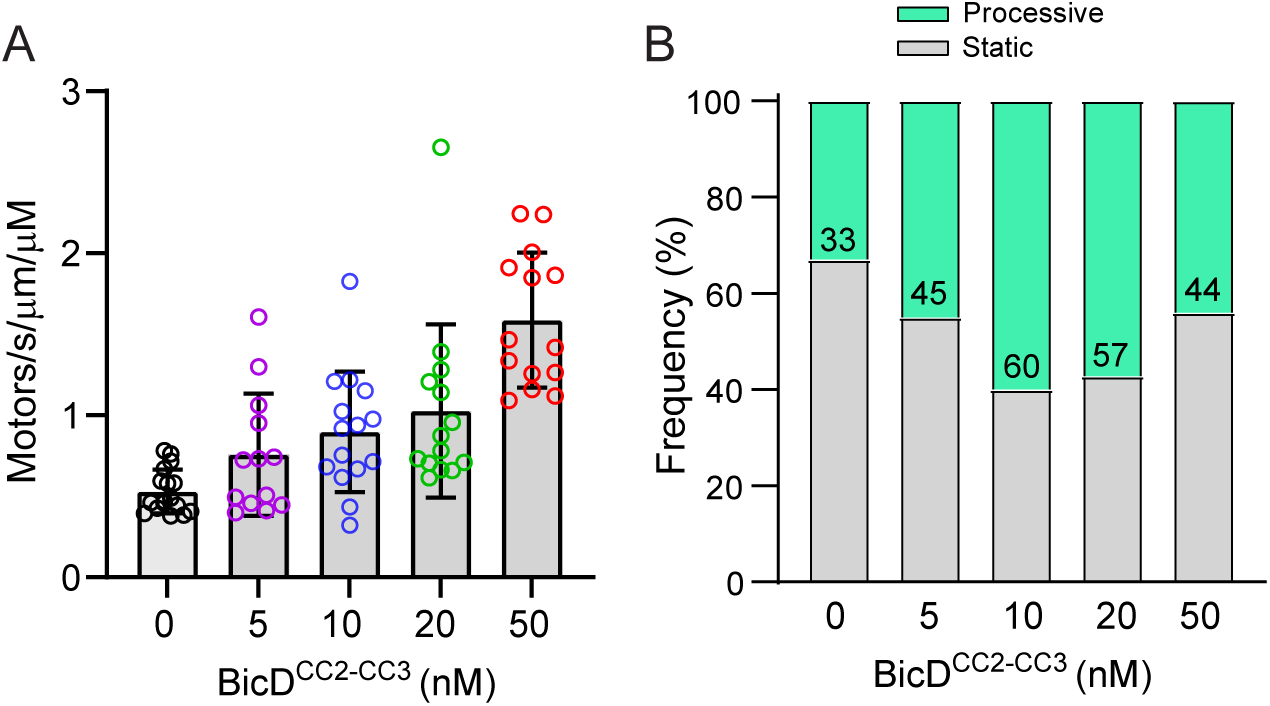
Effect of BicD concentration on the recruitment of kinesin to the microtubule and on kinesin processivity. ***A***, Recruitment of kinesin to the MT at varying BicD^CC2-CC3^ concentrations. BicD^CC2-CC3^ was labeled with a Qdot, and kinesin was unlabeled. N=1. ***B***, Percentage of bound kinesins that showed processive motion at varying BicD concentration. BicD and kinesin were pre-incubated at 100-fold higher concentration and diluted just prior to visualization.

**Supporting Figure 2.**
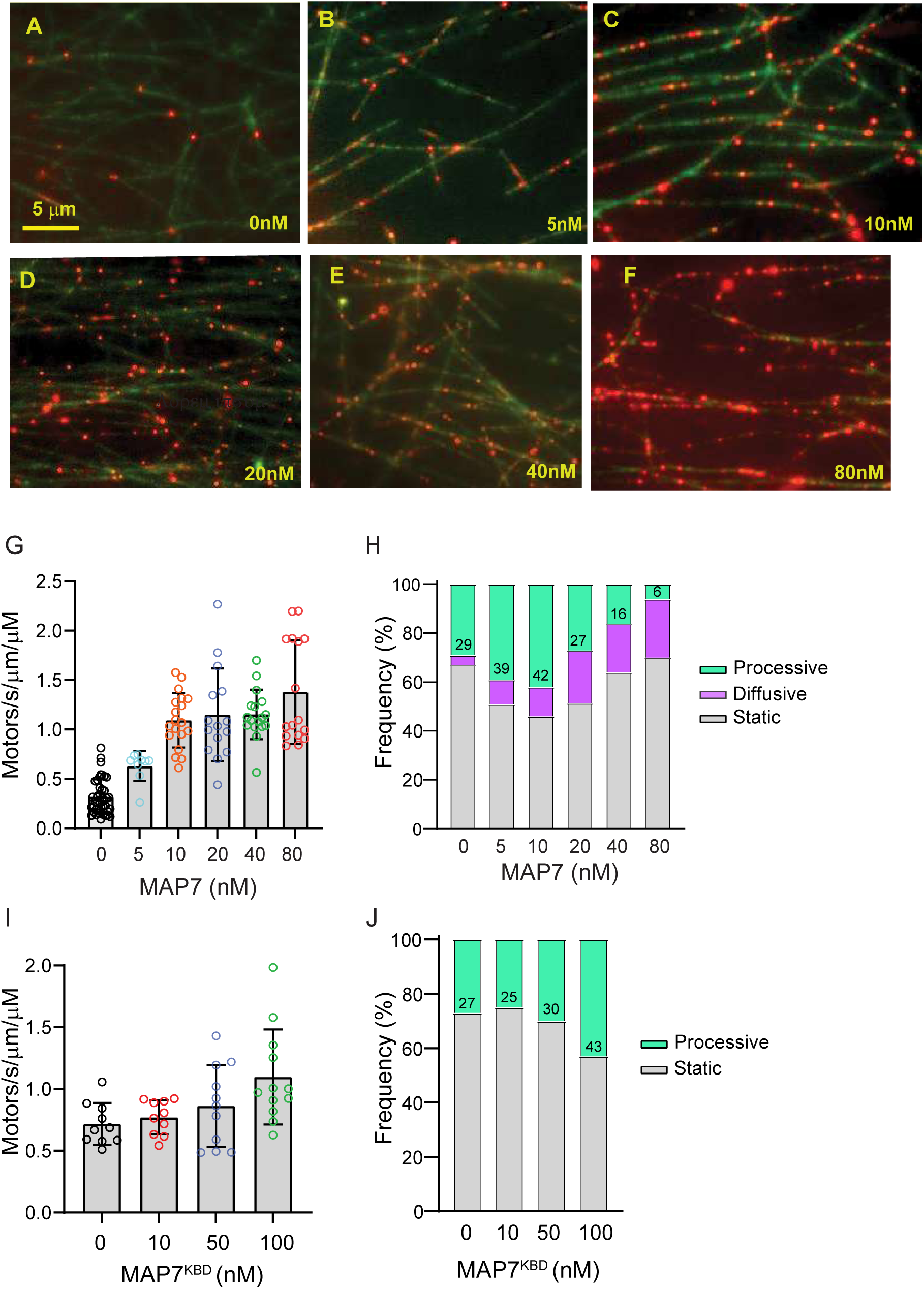
Effect of MAP7 and MAP7^KBD^ concentration on the recruitment of kinesin to the microtubule and on kinesin processivity. ***A-F***, Images showing the recruitment of kinesin (red) to the MT (green) as a function of MAP7 concentration. ***G***, Recruitment of kinesin to the MT as a function of MAP7 concentration N=1. ***H***, Percentage of bound motors that were processive, diffusive or static as a function of MAP7 concentration. ***I***, Recruitment of kinesin to the MT as a function of MAP7^KBD^ concentration, N=1. ***J***, Percentage of bound motors that were processive, diffusive or static as a function of MAP7^KBD^ concentration.

## Notes

### Competing Interest Statement

The authors have declared no competing interest.

## References

1. Mohler, J., and Wieschaus, E. F. (1986) Dominant maternal-effect mutations of Drosophila melanogaster causing the production of double-abdomen embryos Genetics 112, 803–822 10.1093/genetics/112.4.803

2. Olenick, M. A., and Holzbaur, E. L. F. (2019) Dynein activators and adaptors at a glance J Cell Sci 132, 10.1242/jcs.227132

3. McKenney, R. J., Huynh, W., Tanenbaum, M. E., Bhabha, G., and Vale, R. D. (2014) Activation of cytoplasmic dynein motility by dynactin-cargo adapter complexes Science 345, 337–341 10.1126/science.1254198

4. Schlager, M. A., Hoang, H. T., Urnavicius, L., Bullock, S. L., and Carter, A. P. (2014) In vitro reconstitution of a highly processive recombinant human dynein complex EMBO J 33, 1855–1868 10.15252/embj.201488792

5. Gama, J. B., Pereira, C., Simoes, P. A., Celestino, R., Reis, R. M., Barbosa, D. J. et al. (2017) Molecular mechanism of dynein recruitment to kinetochores by the Rod-Zw10-Zwilch complex and Spindly J Cell Biol 216, 943–960 10.1083/jcb.201610108

6. Lee, I. G., Cason, S. E., Alqassim, S. S., Holzbaur, E. L. F., and Dominguez, R. (2020) A tunable LIC1-adaptor interaction modulates dynein activity in a cargo-specific manner Nat Commun 11, 5695 10.1038/s41467-020-19538-7

7. Dienstbier, M., Boehl, F., Li, X., and Bullock, S. L. (2009) Egalitarian is a selective RNA-binding protein linking mRNA localization signals to the dynein motor Genes Dev 23, 1546–1558 10.1101/gad.531009

8. Januschke, J., Nicolas, E., Compagnon, J., Formstecher, E., Goud, B., and Guichet, A. (2007) Rab6 and the secretory pathway affect oocyte polarity in Drosophila Development 134, 3419–3425 10.1242/dev.008078

9. Liu, Y., Salter, H. K., Holding, A. N., Johnson, C. M., Stephens, E., Lukavsky, P. J. et al. (2013) Bicaudal-D uses a parallel, homodimeric coiled coil with heterotypic registry to coordinate recruitment of cargos to dynein Genes Dev 27, 1233–1246 10.1101/gad.212381.112

10. Splinter, D., Tanenbaum, M. E., Lindqvist, A., Jaarsma, D., Flotho, A., Yu, K. L., et al. (2010) Bicaudal D2, dynein, and kinesin-1 associate with nuclear pore complexes and regulate centrosome and nuclear positioning during mitotic entry PLoS Biol 8, e1000350 10.1371/journal.pbio.1000350

11. Larsen, K. S., Xu, J., Cermelli, S., Shu, Z., and Gross, S. P. (2008) BicaudalD actively regulates microtubule motor activity in lipid droplet transport PLoS One 3, e3763 10.1371/journal.pone.0003763

12. McClintock, M. A., Dix, C. I., Johnson, C. M., McLaughlin, S. H., Maizels, R. J., Hoang, H. T., et al. (2018) RNA-directed activation of cytoplasmic dynein-1 in reconstituted transport RNPs Elife 7, 10.7554/eLife.36312

13. Sladewski, T. E., Billington, N., Ali, M. Y., Bookwalter, C. S., Lu, H., Krementsova, E. B., et al. (2018) Recruitment of two dyneins to an mRNA-dependent Bicaudal D transport complex Elife 7, 10.7554/eLife.36306

14. Fagiewicz, R., Crucifix, C., Klos, T., Deville, C., Kieffer, B., Nomine, Y. et al. (2022) In vitro characterization of the full-length human dynein-1 cargo adaptor BicD2 Structure 30, 1470–1478 e1473 10.1016/j.str.2022.08.009

15. Gallisa-Sune, N., Sanchez-Fernandez-de-Landa, P., Zimmermann, F., Serna, M., Regue, L., Paz, J., et al. (2023) BICD2 phosphorylation regulates dynein function and centrosome separation in G2 and M Nat Commun 14, 2434 10.1038/s41467-023-38116-1

16. Zhang, K., Foster, H. E., Rondelet, A., Lacey, S. E., Bahi-Buisson, N., Bird, A. W. et al. (2017) Cryo-EM Reveals How Human Cytoplasmic Dynein Is Auto-inhibited and Activated Cell 169, 1303–1314 e1318 10.1016/j.cell.2017.05.025

17. Urnavicius, L., Lau, C. K., Elshenawy, M. M., Morales-Rios, E., Motz, C., Yildiz, A. et al. (2018) Cryo-EM shows how dynactin recruits two dyneins for faster movement Nature 554, 202–206 10.1038/nature25462

18. Urnavicius, L., Zhang, K., Diamant, A. G., Motz, C., Schlager, M. A., Yu, M., et al. (2015) The structure of the dynactin complex and its interaction with dynein Science 347, 1441-1446 10.1126/science.aaa4080

19. Chowdhury, S., Ketcham, S. A., Schroer, T. A., and Lander, G. C. (2015) Structural organization of the dynein-dynactin complex bound to microtubules Nat Struct Mol Biol 22, 345–347 10.1038/nsmb.2996

20. Canty, J. T., and Yildiz, A. (2020) Activation and Regulation of Cytoplasmic Dynein Trends Biochem Sci 45, 440–453 10.1016/j.tibs.2020.02.002

21. Cai, D., Hoppe, A. D., Swanson, J. A., and Verhey, K. J. (2007) Kinesin-1 structural organization and conformational changes revealed by FRET stoichiometry in live cells J Cell Biol 176, 51–63 10.1083/jcb.200605097

22. Coy, D. L., Hancock, W. O., Wagenbach, M., and Howard, J. (1999) Kinesin’s tail domain is an inhibitory regulator of the motor domain Nat Cell Biol 1, 288–292 10.1038/13001

23. Dietrich, K. A., Sindelar, C. V., Brewer, P. D., Downing, K. H., Cremo, C. R., and Rice, S. E. (2008) The kinesin-1 motor protein is regulated by a direct interaction of its head and tail Proc Natl Acad Sci U S A 105, 8938–8943 10.1073/pnas.0803575105

24. Friedman, D. S., and Vale, R. D. (1999) Single-molecule analysis of kinesin motility reveals regulation by the cargo-binding tail domain Nat Cell Biol 1, 293–297 10.1038/13008

25. Kaan, H. Y., Hackney, D. D., and Kozielski, F. (2011) The structure of the kinesin-1 motor-tail complex reveals the mechanism of autoinhibition Science 333, 883–885 10.1126/science.1204824

26. Stock, M. F., Guerrero, J., Cobb, B., Eggers, C. T., Huang, T. G., Li, X., et al. (1999) Formation of the compact confomer of kinesin requires a COOH-terminal heavy chain domain and inhibits microtubule-stimulated ATPase activity J Biol Chem 274, 14617–14623 10.1074/jbc.274.21.14617

27. Tan, Z., Yue, Y., Leprevost, F., Haynes, S., Basrur, V., Nesvizhskii, A. I., et al. (2023) Autoinhibited kinesin-1 adopts a hierarchical folding pattern Elife 12, 10.7554/eLife.86776

28. Weijman, J. F., Yadav, S. K. N., Surridge, K. J., Cross, J. A., Borucu, U., Mantell, J. et al. (2022) Molecular architecture of the autoinhibited kinesin-1 lambda particle Sci Adv 8, eabp9660 10.1126/sciadv.abp9660

29. Chiba, K., Ori-McKenney, K. M., Niwa, S., and McKenney, R. J. (2022) Synergistic autoinhibition and activation mechanisms control kinesin-1 motor activity Cell Rep 39, 111016 10.1016/j.celrep.2022.111016

30. Barlan, K., Lu, W., and Gelfand, V. I. (2013) The microtubule-binding protein ensconsin is an essential cofactor of kinesin-1 Curr Biol 23, 317–322 10.1016/j.cub.2013.01.008

31. Hooikaas, P. J., Martin, M., Muhlethaler, T., Kuijntjes, G. J., Peeters, C. A. E., Katrukha, E. A., et al. (2019) MAP7 family proteins regulate kinesin-1 recruitment and activation J Cell Biol 218, 1298–1318 10.1083/jcb.201808065

32. Metivier, M., Monroy, B. Y., Gallaud, E., Caous, R., Pascal, A., Richard-Parpaillon, L., et al. (2019) Dual control of Kinesin-1 recruitment to microtubules by Ensconsin in Drosophila neuroblasts and oocytes Development 146, 10.1242/dev.171579

33. Sung, H. H., Telley, I. A., Papadaki, P., Ephrussi, A., Surrey, T., and Rorth, P. (2008) Drosophila ensconsin promotes productive recruitment of Kinesin-1 to microtubules Dev Cell 15, 866–876 10.1016/j.devcel.2008.10.006

34. Chaudhary, A. R., Lu, H., Krementsova, E. B., Bookwalter, C. S., Trybus, K. M., and Hendricks, A. G. (2019) MAP7 regulates organelle transport by recruiting kinesin-1 to microtubules J Biol Chem 294, 10160–10171 10.1074/jbc.RA119.008052

35. Ferro, L. S., Fang, Q., Eshun-Wilson, L., Fernandes, J., Jack, A., Farrell, D. P. et al. (2022) Structural and functional insight into regulation of kinesin-1 by microtubule-associated protein MAP7 Science 375, 326–331 10.1126/science.abf6154

36. Furuta, K., Furuta, A., Toyoshima, Y. Y., Amino, M., Oiwa, K., and Kojima, H. (2013) Measuring collective transport by defined numbers of processive and nonprocessive kinesin motors Proc Natl Acad Sci U S A 110, 501–506 10.1073/pnas.1201390110

37. Young, G., Hundt, N., Cole, D., Fineberg, A., Andrecka, J., Tyler, A. et al. (2018) Quantitative mass imaging of single biological macromolecules Science 360, 423–427 10.1126/science.aar5839

38. Grigoriev, I., Splinter, D., Keijzer, N., Wulf, P. S., Demmers, J., Ohtsuka, T. et al. (2007) Rab6 regulates transport and targeting of exocytotic carriers Dev Cell 13, 305–314 10.1016/j.devcel.2007.06.010

39. Fenton, A. R., Jongens, T. A., and Holzbaur, E. L. F. (2021) Mitochondrial adaptor TRAK2 activates and functionally links opposing kinesin and dynein motors Nat Commun 12, 4578 10.1038/s41467-021-24862-7

40. Canty, J. T., Hensley, A., Aslan, M., Jack, A., and Yildiz, A. (2023) TRAK adaptors regulate the recruitment and activation of dynein and kinesin in mitochondrial transport Nat Commun 14, 1376 10.1038/s41467-023-36945-8

41. Kendrick, A. A., Dickey, A. M., Redwine, W. B., Tran, P. T., Vaites, L. P., Dzieciatkowska, M. et al. (2019) Hook3 is a scaffold for the opposite-polarity microtubule-based motors cytoplasmic dynein-1 and KIF1C J Cell Biol 218, 2982–3001 10.1083/jcb.201812170

42. Siddiqui, N., Zwetsloot, A. J., Bachmann, A., Roth, D., Hussain, H., Brandt, J., et al. (2019) PTPN21 and Hook3 relieve KIF1C autoinhibition and activate intracellular transport Nat Commun 10, 2693 10.1038/s41467-019-10644-9

43. Cronan, J. E., Jr. (1990) Biotination of proteins in vivo. A post-translational modification to label, purify, and study proteins J Biol Chem 265, 10327–10333, https://www.ncbi.nlm.nih.gov/pubmed/2113052

